# Bayesian spatial prediction of livestock tick abundance to support surveillance in data-sparse regions

**DOI:** 10.64898/2026.04.16.719086

**Authors:** Abrar Hussain, Sabir Hussain, Lelys Bravo de Guenni, Rebecca L. Smith

## Abstract

Fragmented and geographically uneven surveillance limits the ability to target tick monitoring and control in many livestock systems. We evaluated whether heterogeneous published surveillance could be converted into district-level, uncertainty-aware abundance predictions for *Rhipicephalus microplus* and *Hyalomma anatolicum* across Punjab and Khyber Pakhtunkhwa, Pakistan. Species-specific tick counts were standardized by the number of animals examined and sampling duration and expressed as ticks per 100,000 animal-months. Environmental covariates were summarized using principal component analysis, with livestock density included separately. Bayesian Gaussian models, with and without a district-level Besag-York-Mollie 2 spatial effect, were fitted in R using Integrated Nested Laplace Approximation (INLA) and compared. Leave-one-district-out validation was performed to assess out-of-sample predictive performance. Spatial models improved model fit for both species. For *R. microplus*, PC1 and PC4 showed credible negative associations with abundance, whereas no fixed-effect 95% credible interval excluded zero for *H. anatolicum*. Predicted *R. microplus* abundance was more spatially heterogeneous, with high but uncertain estimates in several northern districts, whereas *H. anatolicum* predictions were more homogeneous and relatively higher toward southern Punjab. Mapping posterior uncertainty identified districts where additional standardized surveillance would be most informative. This approach provides a transferable framework for extracting spatial surveillance information from heterogeneous veterinary data while explicitly representing uncertainty.

## 1. Introduction

Animal-health surveillance provides the evidence needed to guide disease prevention and control, but its usefulness depends on the availability of comparable and geographically informative data (Hoinville et al., 2013). In data-sparse settings, surveillance information is often derived from studies conducted for different purposes, in different locations, over different periods, and with varying sampling intensities, limiting direct comparisons across areas. Spatially explicit approaches that account for sampling heterogeneity and quantify uncertainty can help extract additional information from such fragmented evidence, generate more comparable estimates across locations, and identify where further surveillance may be most valuable (Reist et al., 2012). This challenge is particularly relevant to livestock ticks, for which surveillance is often geographically incomplete despite their substantial animal health and economic importance.

Among livestock health threats requiring effective surveillance, ticks and tick-borne diseases impose substantial economic and animal-health burdens worldwide, with estimated annual losses of US$13.9-18.7 billion in cattle production and associated by-products (Karim et al., 2017). The challenge is particularly important in Pakistan, where livestock contributes more than 60% of agricultural value added and about 11% of national GDP, and approximately 35 million rural residents depend on the sector for their livelihoods (Rajput et al., 2023; Rehman et al., 2017a). Within this production system, ticks compromise animal health and productivity by blood-feeding and by transmitting a wide range of pathogens that affect both veterinary and human health (Dantas-Torres et al., 2012). In Pakistan, multiple tick species infest bovines and small ruminants, with *Rhipicephalus microplus* and *Hyalomma anatolicum* among the most frequently reported species from domestic livestock (Hussain et al., 2021; Jabbar et al., 2015; Rehman et al., 2019, 2017b). *R. microplus* (the cattle tick) transmits agents of bovine tick fever, including *Anaplasma marginale, Babesia bigemina*, and *B. bovis*, whereas *H. anatolicum* transmits *Theileria annulata* and Crimean-Congo hemorrhagic fever virus (Ahmed et al., 2021; Jabbar et al., 2015; Mourya et al., 2019). Human exposure to zoonotic tick-borne pathogens can occur through bites during herd management, manual tick removal, or contact with blood and tissues from infected animals (Ahmed et al., 2021).

Tick occurrence and abundance vary spatially in response to climate, landscape, and host availability, making these factors relevant to spatial surveillance. Temperature, precipitation, relative humidity, and elevation influence tick development, survival, and habitat suitability, while livestock availability can affect opportunities for host-dependent population maintenance (Bonnet et al., 2022; De Clercq et al., 2012; Rajput et al., 2023). These relationships may differ between species because *R. microplus* is a one-host tick closely associated with cattle, whereas *H. anatolicum* can use multiple hosts across its life cycle (Bonnet et al., 2022; Vatansever, 2018). Incorporating environmental and livestock information may therefore improve spatial inference where direct surveillance observations are sparse.

Despite their veterinary importance and the environmental heterogeneity likely to shape their abundance, spatially comparable estimates of *R. microplus* and *H. anatolicum* abundance remain limited in Pakistan. A recent systematic review synthesized published district-level records of these species across Pakistan and demonstrated substantial geographic heterogeneity in their reported distributions (Hussain et al., 2024). However, the underlying studies differed substantially in the number of animals examined, sampling duration, and geographic coverage, limiting direct comparison of tick burden among districts. The previous synthesis was therefore useful for describing where the species had been reported, but it did not standardize observations by sampling effort, generate model-based abundance predictions for unsampled districts, or quantify district-level prediction uncertainty. These limitations restrict the use of the existing evidence for geographically targeted surveillance planning.

We therefore asked whether heterogeneous and geographically incomplete published tick-surveillance records could be standardized and integrated with environmental, livestock, and spatial information to predict district-level abundance and identify surveillance gaps. Specifically, we aimed to (1) derive comparable district-level abundance estimates while accounting for differences in sampling effort; and (2) generate uncertainty-aware spatial predictions for unsampled districts, evaluate out-of-sample predictive performance, and identify areas where additional standardized surveillance would be most informative. This framework could help target limited surveillance resources and be transferable to other data-sparse veterinary settings where surveillance data are fragmented or incomplete.

## 2. Materials and Methods

### 2.1 Study area and data curation

This study focuses on Punjab and Khyber Pakhtunkhwa (KP), the two major provinces of Pakistan, which together host over 60% of the national livestock population, with approximately 104 million in Punjab and 48.7 million in KP (MSN, 2025). Districts were used as the spatial units of analysis. Tick data were derived from the PRISMA-guided systematic review (Hussain et al., 2024), which compiled district-level records of *R. microplus* and *H. anatolicum* from published studies in Pakistan from 1990 to 2023. For the present analysis, the underlying source studies were revisited to verify species-specific tick counts, sampling duration, and the number of animals examined. The analysis was restricted to Punjab and KP because these provinces provided substantially greater district-level data coverage, whereas records from other provinces were too sparse for reliable spatial prediction. For prediction purposes, districts without usable species-specific abundance observations were considered unsampled. For each sampled district, exposure-adjusted tick abundance was calculated as:

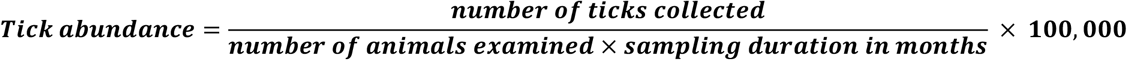

The resulting measure represents the number of ticks collected per 100,000 animal-months of sampling effort and accounts for differences in both the number of animals examined and sampling duration. All included studies collected ticks directly from host animals by hand-picking; no environmental trapping methods were represented. District-level administrative boundaries were obtained as polygon shapefiles for spatial analysis and mapping (Admin Boundaries, 2024). The two tick species were analyzed separately using the same analytical workflow, summarized in Figure 1.

**Figure 1.**
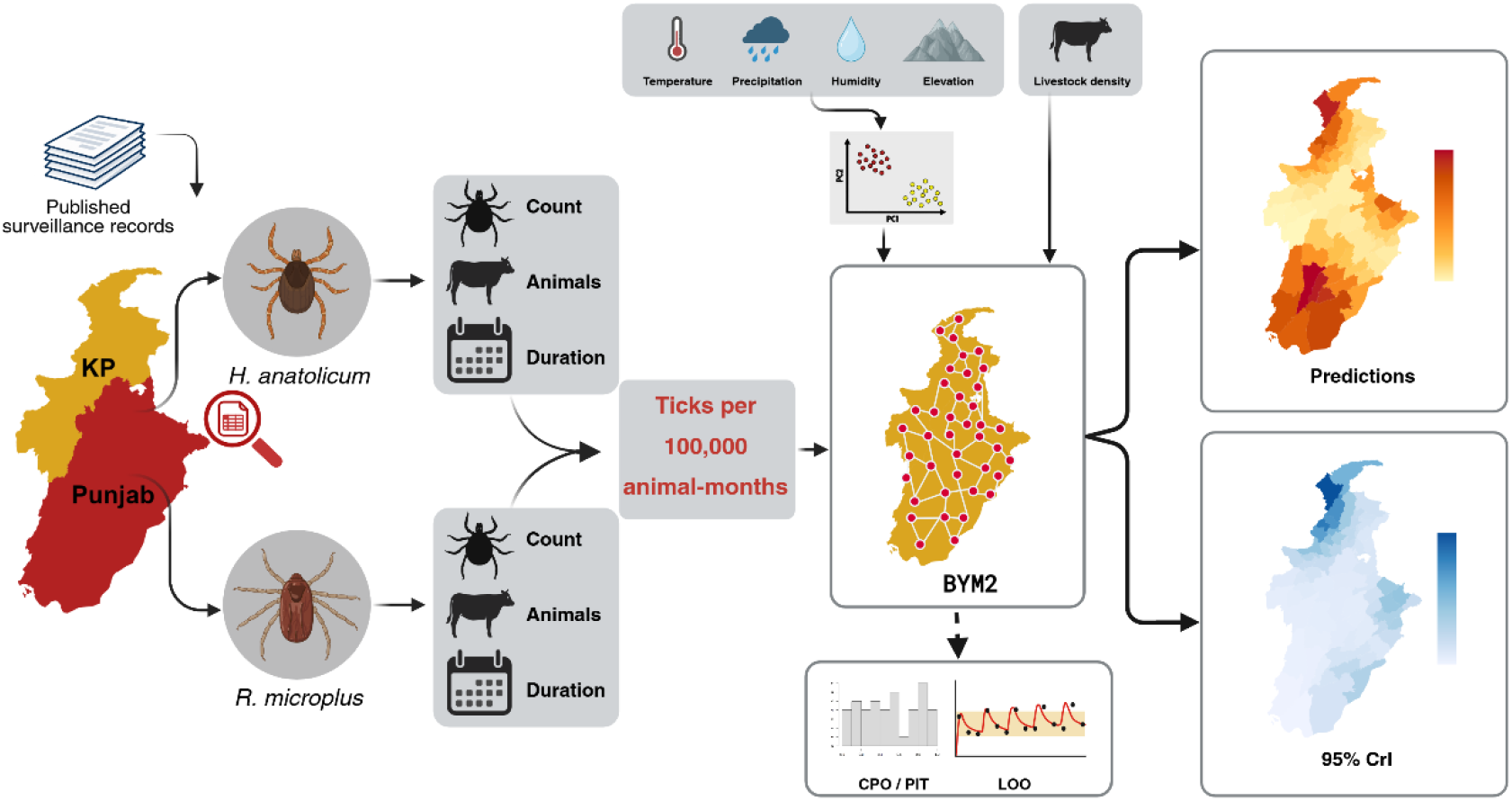
Overview of the analytical workflow used to generate district-level Bayesian spatial predictions of *Rhipicephalus microplus* and *Hyalomma anatolicum* abundance in Punjab and Khyber Pakhtunkhwa, Pakistan. (Created with BioRender.com).

### 2.2 Environmental and host covariates

Mean annual temperature and annual precipitation were obtained from the WorldClim version 2.1 at a spatial resolution of 2.5 arcminutes (corresponding to approximately 5 × 5 km grid cells), and mean values were extracted for each district polygon (WorldClim, 2025). Elevation data were derived with a zoom level of 7 (approximately 600 m), and district-level mean elevation was calculated. Relative humidity at 2 m above the surface (RH2M) was obtained from NASA POWER for the 1990-2023 period. Annual relative humidity values were retrieved on a 1° x 1° grid across Pakistan, spatially interpolated using inverse distance weighting (IDW) to a 0.5° grid, and the mean interpolated relative humidity was extracted for each district polygon (NASA, 2025). District-level livestock population data were obtained from the Pakistan Livestock Census 2006 (PBS, 2006). Cattle, buffaloes, sheep, goats, horses, mules, asses, and camels were summed to obtain the total livestock population for each district. Livestock census data were harmonized with the district boundaries used in the spatial analysis before calculating district-level livestock density, obtained by dividing the total livestock population by district area and expressed as animals per km^2^. The 2006 census was used to represent broad spatial variation in livestock host availability during the period represented by the tick surveillance records.

Relationships among the four environmental covariates were assessed using Pearson correlation coefficients and visualized using a correlation heatmap. Because substantial correlations were present among the environmental variables, principal component analysis (PCA) was used to transform the correlated environmental covariates into mutually orthogonal components and reduce multicollinearity among predictors. Livestock density was retained as a separate host covariate and was not included in the environmental PCA.

### 2.3 Principal component analysis (PCA)

PCA was performed on district-level mean temperature, precipitation, relative humidity, and elevation using the *prcomp()* function in **R** after centering and scaling the variables, thereby generating mutually orthogonal environmental components(Abdi and Williams, 2010). Scores for PC1-PC4 were extracted and joined to the spatial dataset. The proportion of variance explained by each component was examined using a scree plot. A PCA biplot was also used to visualize district distributions along the first two principal components and the directions and magnitudes of environmental variable loadings, with districts distinguished by province.

### 2.4 Outcome and covariate preparation

Prior to Gaussian modeling, the distributions of the species-specific outcome variables were evaluated using histograms, quantile-quantile plots, and Shapiro-Wilk tests. To improve numerical stability during model fitting, abundance rates for both species were first expressed in thousands of ticks per 100,000 animal-months by dividing the rates by 1,000. The rescaled *R. microplus* abundance rate remained substantially right-skewed and was therefore natural-log-transformed prior to modeling, whereas the rescaled *H. anatolicum* abundance rate was approximately Gaussian and was retained without further transformation (Figure S1). Livestock density showed only mild skewness and was standardized to a mean of zero and a standard deviation of one prior to modeling to improve numerical stability and allow its regression coefficient to be interpreted as the change in the outcome associated with a one-standard-deviation increase in livestock density (Figure S2).

### 2.5 Statistical framework

A Bayesian modeling framework was used to estimate associations between environmental and host covariates and district-level tick abundance. Gaussian likelihoods were used for both species, with PC1-PC4 and standardized livestock density included as fixed effects. For each tick species, two model formulations were fitted: (i) a nonspatial Gaussian regression model containing only the fixed effects and (ii) a spatial Gaussian model containing the same fixed effects together with a district-level Besag-York-Mollie 2 (BYM2) random effect (Quick et al., 2021). The BYM2 formulation accounts for residual district-level variation through both spatially structured and unstructured components. For each tick species, spatial and nonspatial models were compared using the Deviance Information Criterion (DIC) and Watanabe-Akaike Information Criterion (WAIC), with lower values indicating better relative fit.

Spatial dependence was defined using Queen contiguity, in which districts sharing a boundary or a vertex were considered neighbors. The resulting adjacency network was checked for isolated districts and overall connectivity before conversion to the Integrated Nested Laplace Approximation (INLA) graph format. Bayesian models were fitted in R using the Integrated INLA framework with default priors (Lindgren and Rue, 2015; Rue et al., 2009). The spatial Gaussian BYM2 model for the district *i*was specified as:

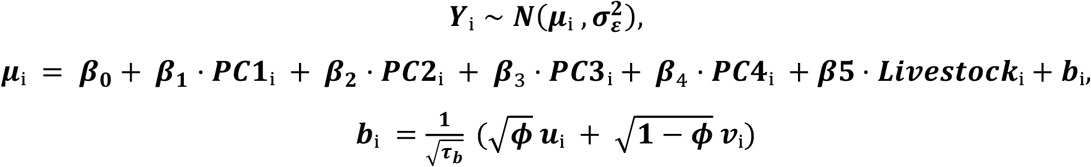

where ***Y***_***i***_is the modeled tick abundance rate for district ***i***, corresponding to log(*R*/1000)for *R. microplus* and *H*/1000 for *H. anatolicum*; ***μ***_***i***_is the expected value of the outcome on the corresponding modeling scale; ***β***_**0**_is the intercept; ***β***_**1**_ **− *β***_**4**_ are the regression coefficients for ***PC1-PC4; β***_**5**_is the coefficient for standardized livestock density; and ***σ***^**2**^is the Gaussian residual variance. The term ***b***_***i***_represents the district-level BYM2 random effect, ***τ***_***b***_controls its overall precision, and ***ϕ*** is the mixing parameter representing the proportion of random-effect variance attributable to the spatially structured component. The terms ***u***_***i***_and *v*_*i*_represent the structured spatial and unstructured district-specific components, respectively. For the nonspatial model, ***b***_***i***_was omitted.

### 2.6 District-level prediction and leave-one-out validation

Posterior predictions from the final spatial BYM2 models were generated for all districts in Punjab and KP. District-level posterior medians and corresponding 95% credible intervals (CrIs) were extracted from the fitted posterior distributions. Because *R. microplus* was modeled as log(*R*/1000), posterior predictions and credible limits were exponentiated and multiplied by 1,000 to return them to the original abundance-rate scale. For *H. anatolicum*, which was modeled as *H*/1000, posterior predictions and credible limits were multiplied by 1,000. Final estimates for both species were therefore expressed as predicted ticks per 100,000 animal-months and mapped at the district level. To characterize spatial variation in prediction precision, district-level 95% CrI widths were mapped, with wider intervals indicating greater predictive uncertainty and areas where additional standardized surveillance could be most informative.

Predictive performance and calibration were assessed using conditional predictive ordinate (CPO) and probability integral transform (PIT) diagnostics. In addition, leave-one-district-out crossvalidation was performed separately for each species across districts with observed abundance data to evaluate out-of-sample predictive performance. In each iteration, the observed outcome for one district was withheld, the spatial BYM2 model was refitted using the remaining observed districts, and abundance in the withheld district was predicted using its environmental covariates, livestock density, and spatial structure. This procedure was repeated until every observed district had an out-of-sample prediction. For each withheld district, the posterior median prediction and its 95% CrI were extracted and back-transformed to the original abundance-rate scale. Validation was summarized as the number and percentage of observed district-level abundance values falling within their corresponding 95% CrIs.

## 3. Results

### 3.1 Descriptive epidemiology of ticks

Observed exposure-adjusted abundance rates of *R. microplus* and *H. anatolicum* showed distinct but partially overlapping spatial patterns across Punjab and KP. For *R. microplus*, abundance data were available for 44 of 71 districts. Sampling effort varied substantially among the underlying studies, with between 184 and 12,546 animals examined per district (mean = 3,912; SD = 3,307). Higher abundance rates, reaching approximately 6,000-7,000 ticks per 100,000 animal-months, occurred in several districts of northern KP as well as central and southern Punjab. Moderate abundance rates were distributed across both provinces, whereas lower rates were observed in several districts of western KP and Punjab (Figure 2). For *H. anatolicum*, abundance data were available for 39 of 71 districts, with a similarly heterogeneous sampling effort (mean = 4,419 animals examined; SD = 3,422). The highest observed abundance rates, exceeding 4,000 ticks per 100,000 animal-months, occurred primarily in southern and central Punjab. Moderate abundance rates of approximately 2,000-3,000 ticks per 100,000 animal-months were distributed broadly across Punjab and parts of KP, whereas lower rates were observed mainly in northwestern districts (Figure 2).

**Figure 2.**
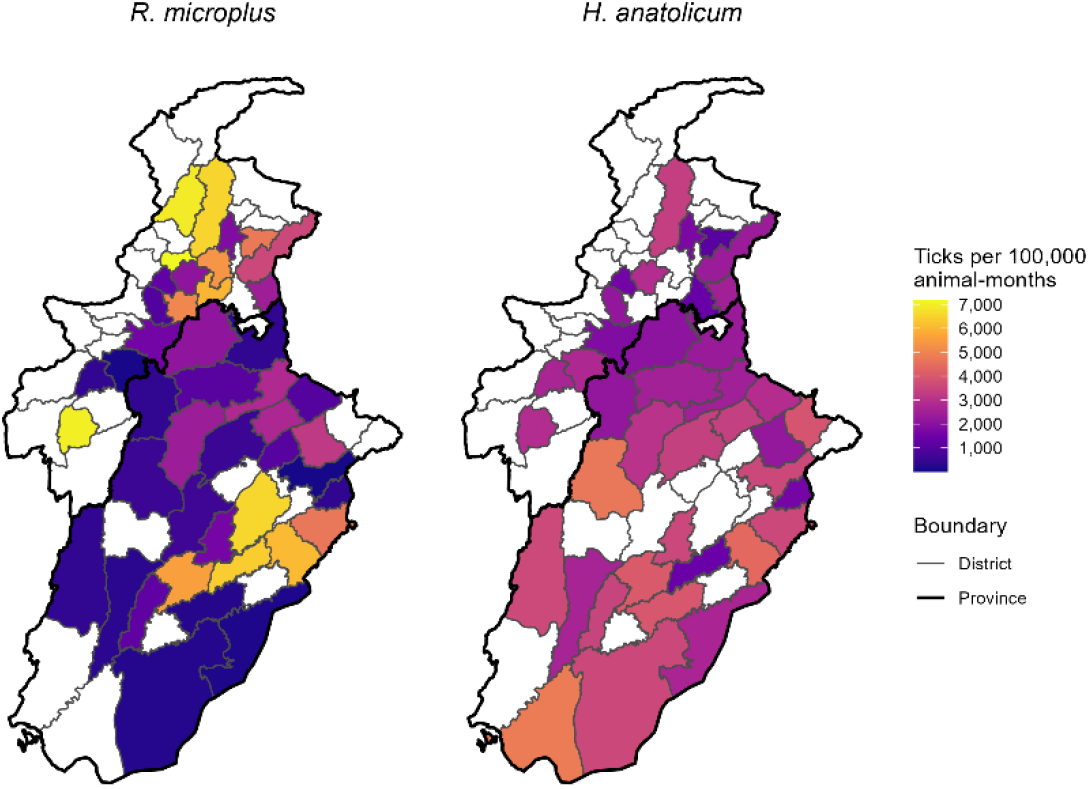
Observed district-level exposure-adjusted abundance of *R. microplus* and *H. anatolicum* in Punjab and Khyber Pakhtunkhwa, Pakistan. Abundance is expressed as ticks per 100,000 animal-months. White districts indicate locations without usable abundance data. *(Map lines delineate study areas and do not necessarily depict accepted national boundaries*.)

### 3.2 Covariates and principal component analysis

The environmental covariates showed substantial spatial variation across the region and were moderately to strongly correlated with one another, supporting the use of PCA to obtain orthogonal environmental predictors (Figure 3; Figure S3). The first two principal components accounted for 98.40% of the total environmental variation, with PC1 explaining 80.82% and PC2 explaining 17.58%. PC3 and PC4 explained only 1.48% and 0.12% of the variance, respectively (Figure S4); however, all four components were retained in the final modeling framework to represent the complete set of orthogonal environmental gradients. PC1 represented a gradient toward warmer, drier, lower-elevation, and lower-humidity conditions, while PC2 was driven primarily by a strong positive loading for precipitation, with positive contributions from temperature and relative humidity, and a negative loading for elevation, broadly representing a wetter, lower-elevation environmental gradient. The remaining two components captured comparatively small residual variation among the environmental variables. District scores on PC1 and PC2 showed geographic separation between portions of Punjab and KP, reflecting the contrasting climatic and topographic conditions across the study area (Figure 4). Livestock density also varied substantially among districts, ranging from approximately 36 to 839 animals/km^2^, with a median of approximately 292 animals/km^2^ (Figure 3).

**Figure 3.**
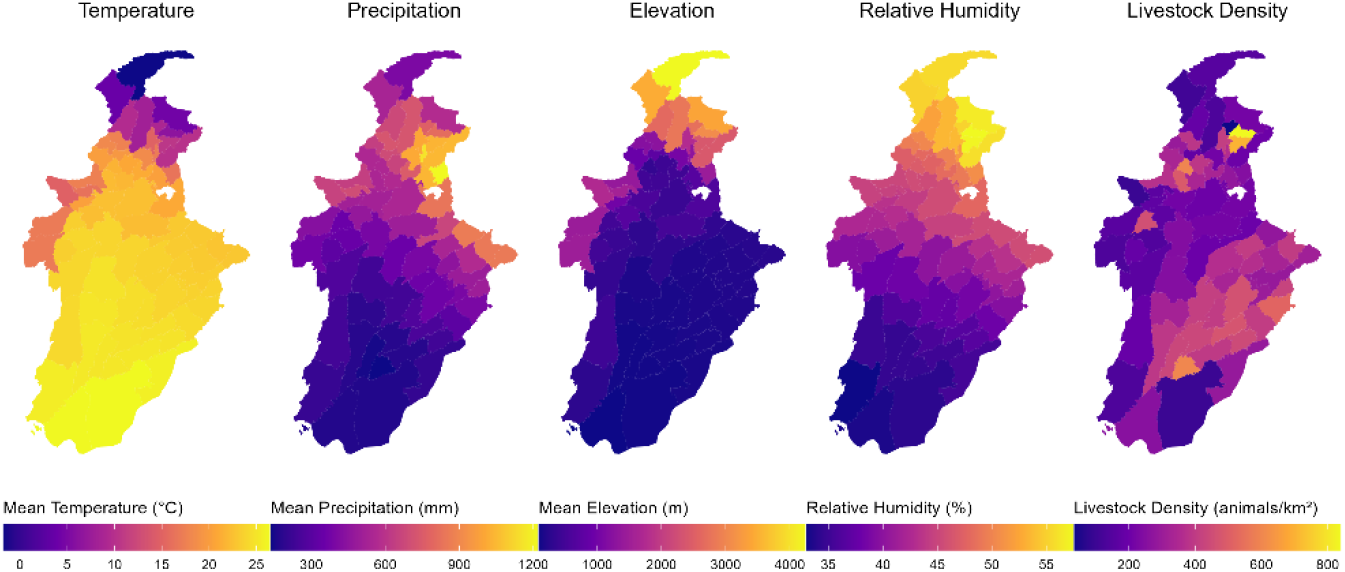
Spatial distribution of district-level environmental and host covariates across Punjab and Khyber Pakhtunkhwa, Pakistan, including mean annual temperature, annual precipitation, mean elevation, relative humidity, and livestock density.

**Figure 4.**
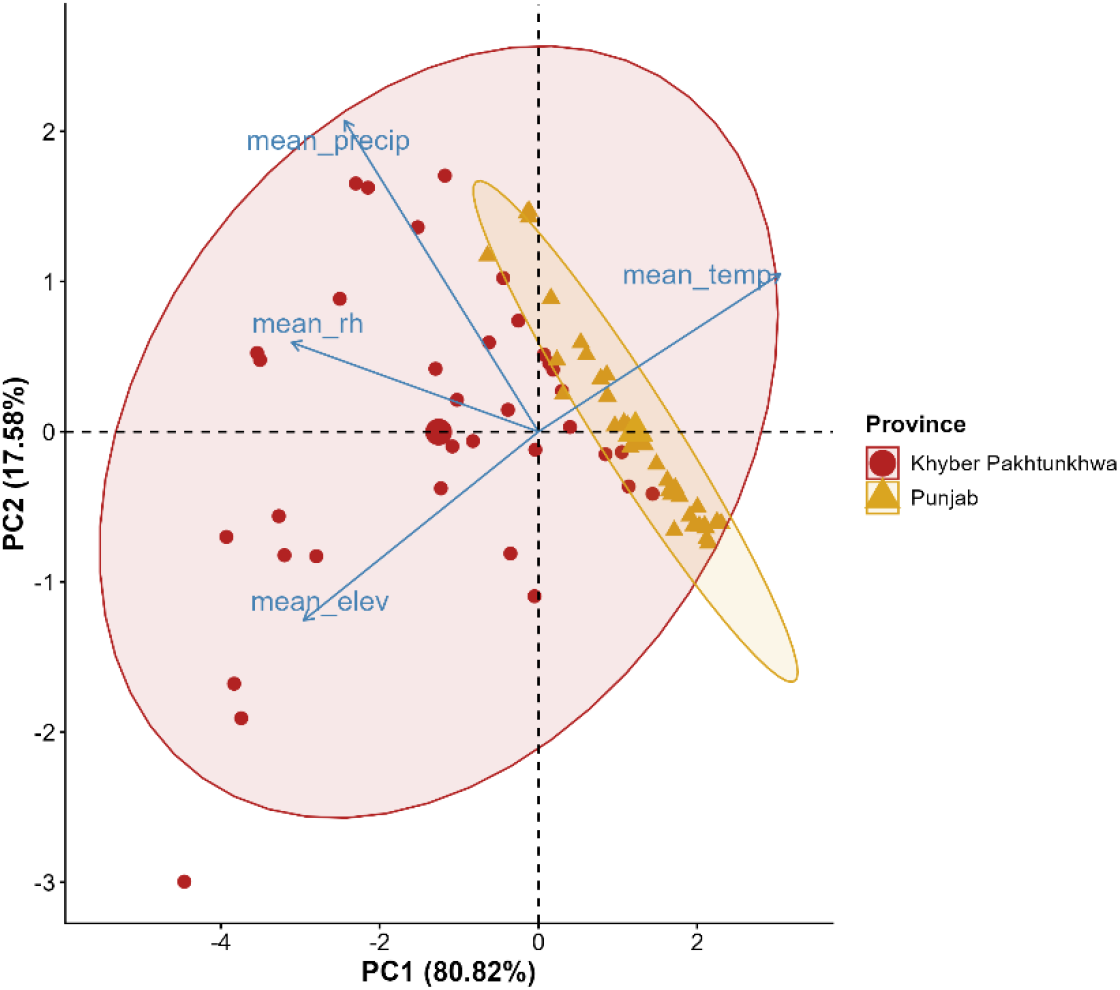
Principal component analysis of district-level environmental covariates across Punjab and Khyber Pakhtunkhwa. The biplot shows district scores on PC1 and PC2, with vectors indicating the direction and relative contribution of mean temperature, precipitation, relative humidity, and elevation to the two principal components. Districts are distinguished by province, with ellipses indicating their distribution within each province.

### 3.3 Bayesian model estimates

For both tick species, inclusion of the district-level BYM2 random effect improved model fit relative to the corresponding nonspatial Gaussian model. The spatial BYM2 models containing PC1-PC4 and standardized livestock density were therefore retained as the final models for inference and prediction (Table S1). Posterior estimates from the final BYM2 models differed between species (Table 1). Only *R. microplus* showed clearly supported fixed-effect associations, with abundance decreasing along PC1 and PC4; PC2, PC3, and livestock density had 95% CrIs spanning zero. For *H. anatolicum*, PC1 showed a positive and PC2 a negative association, but neither was credibly supported because their 95% CrIs included zero; the remaining fixed effects were likewise uncertain.

**Table 1.** Posterior estimates of fixed effects from the final spatial BYM2 models for *Rhipicephalus microplus* and *Hyalomma anatolicum*. Posterior means and 95% credible intervals (CrIs) are shown for the four principal components and standardized livestock density. Bold values indicate predictors whose 95% CrIs do not include zero.

| Species | Predictor | Posterior mean | 95% CrI |
| --- | --- | --- | --- |
| *R. microplus* | PC1 | **−0.47** | −0.77 to −0.15 |
|  | PC2 | −0.40 | −1.14 to 0.34 |
|  | PC3 | −0.03 | −1.71 to 1.67 |
|  | PC4 | **−7.63** | −13.46 to −1.81 |
|  | Livestock density | 0.06 | −0.36 to 0.48 |
| *H. anatolicum* | PC1 | 0.12 | −0.14 to 0.37 |
|  | PC2 | −0.53 | −1.10 to 0.02 |
|  | PC3 | 0.79 | −0.37 to 1.94 |
|  | PC4 | −4.21 | −8.45 to 0.05 |
|  | Livestock density | −0.28 | −0.58 to 0.02 |

### 3.4 Predictions and model validation

The final spatial BYM2 models were used to generate posterior median abundance estimates and 95% credible intervals for all 71 districts, including the unsampled districts. The resulting maps showed substantial spatial heterogeneity in predicted abundance, particularly for *R. microplus*, with several districts in northern KP receiving comparatively high posterior predictions and wide credible intervals. Predicted *H. anatolicum* abundance was more spatially homogeneous across the study region (Figure 5). Spatial variation in predictive uncertainty was also evident, with wider 95% credible intervals particularly in northern KP for *R. microplus* and in northern and western KP for *H. anatolicum*. Under the uncertainty-based surveillance criterion, these districts represented the strongest priorities for additional standardized sampling (Figure 6).

**Figure 5.**
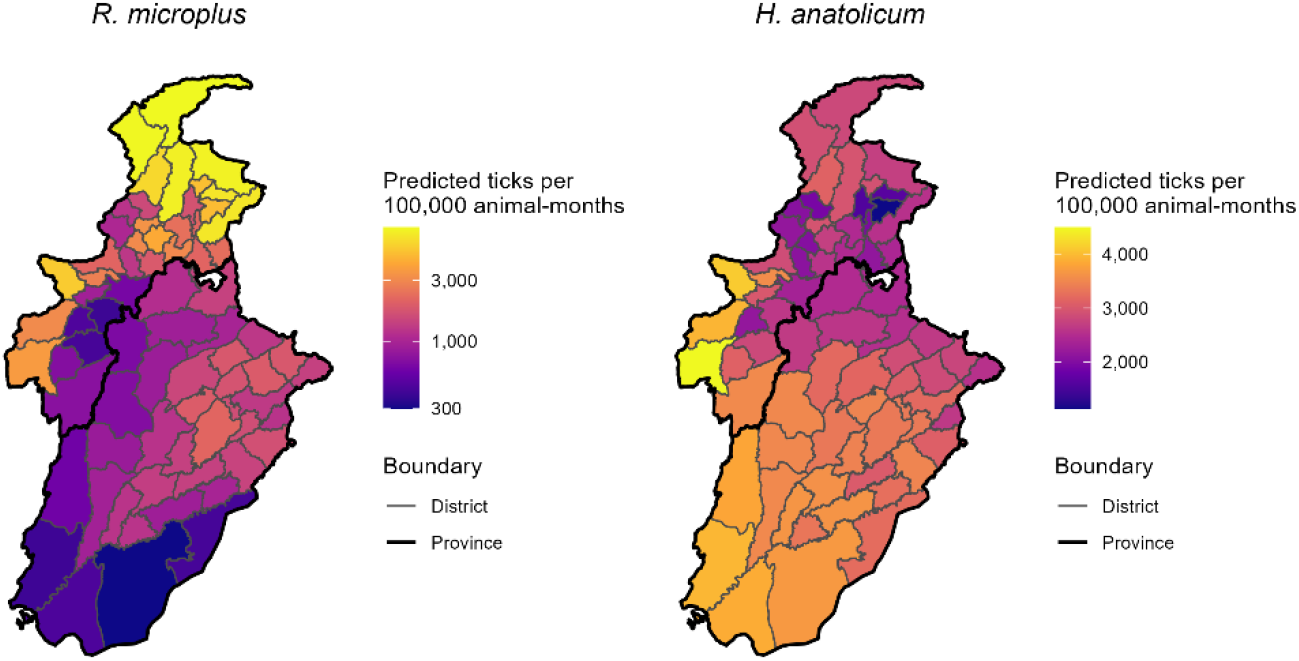
District-level posterior median predictions of exposure-adjusted abundance for *Rhipicephalus microplus* and *Hyalomma anatolicum* across Punjab and Khyber Pakhtunkhwa, Pakistan, based on the final spatial BYM2 models. Abundance is expressed as ticks per 100,000 animal-months. *(Map lines delineate study areas and do not necessarily depict accepted national boundaries*.)

**Figure 6.**
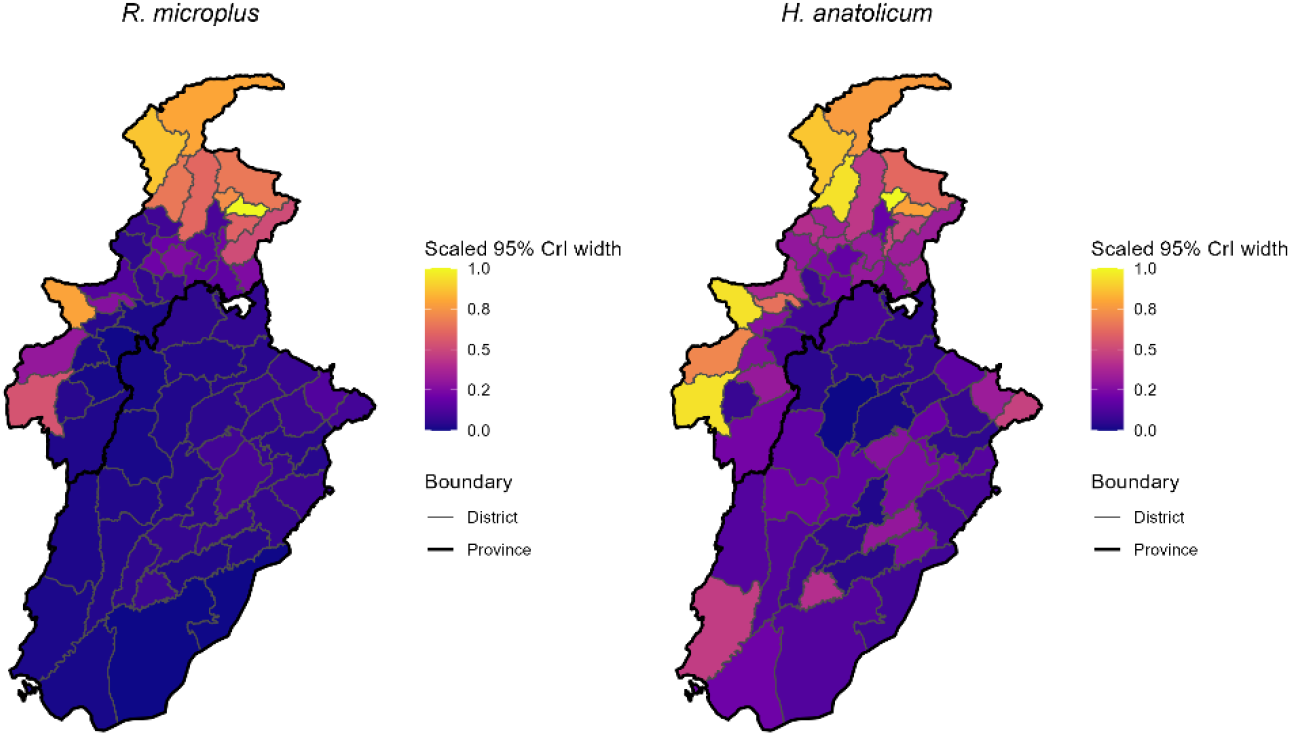
Spatial uncertainty in district-level abundance predictions for *Rhipicephalus microplus* and *Hyalomma anatolicum*, based on 95% credible-interval width. Widths were scaled 0–1 by species; higher values indicate greater uncertainty. *(Map lines delineate study areas and do not necessarily depict accepted national boundaries*.)

Conditional predictive ordinate calculations produced no failures for either species. PIT values were centered near 0.5 for both *R. microplus* (mean = 0.503; median = 0.499) and *H. anatolicum* (mean = 0.506; median = 0.473), indicating balanced predictive calibration across observed districts (Figure S5). Leave-one-district-out validation was used to assess interval inclusion for district-level predictions. Observed abundance fell within the corresponding 95% credible interval for 26 of 44 (59.1%) *R. microplus* districts and 31 of 39 (79.5%) *H. anatolicum* districts (Figure 7; Table S2).

**Figure 7.**
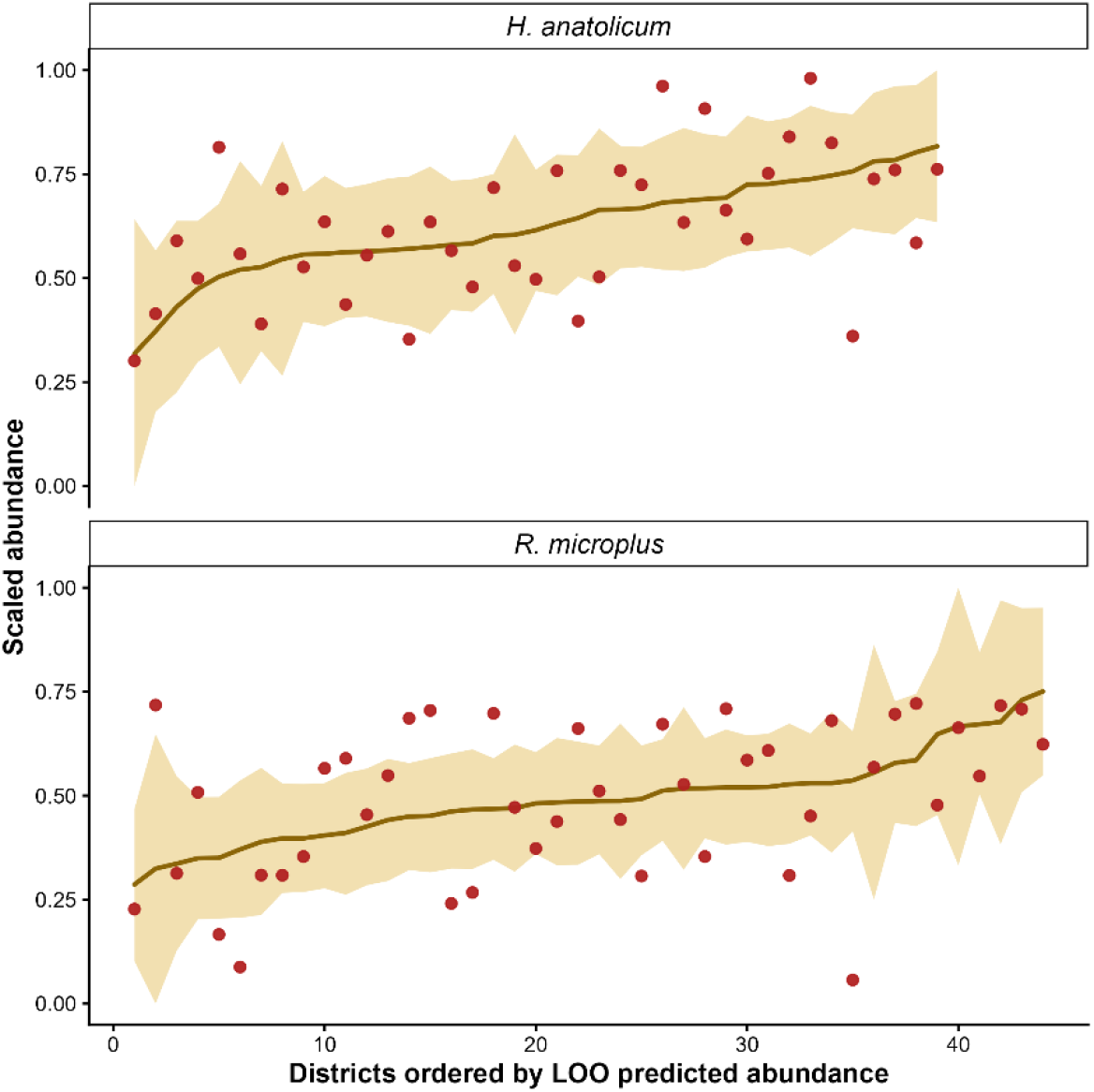
Leave-one-district-out validation of district-level abundance predictions. The line shows LOO posterior median predictions, points show observed abundance, and the shaded band shows 95% credible intervals. Values were scaled separately within species for visualization.

## 4. Discussion

This study demonstrates how heterogeneous published tick-surveillance records can be standardized for differences in sampling effort and integrated within a common spatial framework to generate district-level abundance estimates while explicitly representing uncertainty. Standardizing abundance by the number of animals examined and sampling duration improved comparability among records collected under different sampling intensities, although it could not eliminate all heterogeneity among the underlying studies. The surveillance value of the approach lies not only in identifying areas of comparatively high predicted abundance, but also in distinguishing areas supported by relatively precise information from those where additional standardized sampling would be most informative.

Predicted *R. microplus* abundance was comparatively heterogeneous, with several of the highest posterior estimates occurring in northern KP, whereas *H. anatolicum* showed a more spatially homogeneous pattern with relatively higher abundance toward southern Punjab. Environmental associations also differed between species. For *R. microplus*, PC1 and PC4 showed credible negative associations with abundance. Because higher PC1 scores represented warmer, drier, lower-humidity, and lower-elevation conditions, the negative association indicated greater predicted abundance toward the relatively moister end of this composite gradient, consistent with the sensitivity of *R. microplus* eggs and larvae to desiccation (Sutherst and Bourne, 2006). PC4 was also negatively associated with abundance, but because it accounted for only 0.12% of the environmental variation, this association reflects a comparatively minor residual environmental gradient rather than a dominant environmental pattern. For *H. anatolicum*, PC1 showed a positive and PC2 a negative tendency, although neither association was credibly supported. These directions are broadly consistent with its occurrence in relatively arid environments, but survival of unfed stages can still decline under excessive heat and desiccation (Khogali and Hassan, 2021), indicating that adaptation to arid environments does not eliminate sensitivity to temperature and moisture stress.

District-level livestock density showed no clearly supported association with abundance for either species after accounting for environmental gradients and spatial dependence. This does not imply that host availability is unimportant, but rather that aggregate livestock density may be an incomplete proxy for the host and production conditions relevant to tick populations. For example, similar district-level livestock densities could arise from many small household-based herds or from fewer larger livestock operations, despite potentially substantial differences in animal housing, grazing, movement, veterinary access, biosecurity, and tick-control practices. Such management and production characteristics have been associated with tick infestation in livestock systems in Pakistan (Alvi et al., 2024; Ghafar et al., 2021; Rehman et al., 2017b). Consistent with this interpretation, inclusion of the district-level BYM2 random effect improved model fit for both species, indicating residual geographic variation not explained by the measured environmental and livestock covariates. This residual variation may partly reflect unmeasured differences in host composition, production systems, management practices, landscape connectivity, or other locally structured factors. This remaining geographic variation also makes uncertainty an important part of interpreting the district-level estimates.

Quantifying spatial uncertainty is an important advantage of Bayesian modeling over approaches that provide only point estimates (Cheng et al., 2018). District-level 95% credible interval widths revealed substantial geographic variation in prediction precision (Figure 6). For *R. microplus*, the greatest relative uncertainty was concentrated in northern KP, whereas for *H. anatolicum*, relatively wide credible intervals occurred in northern and western KP. Leave-one-district-out validation provided a complementary out-of-sample assessment, with greater interval inclusion for *H. anatolicum* than for *R. microplus*, indicating more consistent prediction when observed districts were withheld. Together, these results identify areas where additional standardized surveillance would be particularly informative. For *R. microplus*, new sampling in northern KP would help determine whether the comparatively high predicted abundance is supported by field observations, whereas for *H. anatolicum*, additional sampling in northern and western KP would improve prediction precision. Future surveillance would further benefit from consistent recording of host species and numbers examined, sampling duration, season, and month of collection, sampling method, and precise sampling location.

More broadly, integrating observations collected under different study designs within a common exposure-adjusted spatial framework addresses a recurring challenge in veterinary surveillance, where available data often differ in purpose, completeness, geographic coverage, and sampling intensity (Hoinville et al., 2013; Paiba et al., 2007). Bayesian spatial approaches are useful in such settings because they can incorporate spatial dependence, generate predictions for locations with sparse or missing observations, and explicitly represent associated uncertainty (Staubach et al., 2002). Although developed using livestock tick data from Punjab and KP, the same general framework could be applied to other data-sparse veterinary systems where records include location, abundance, and sampling effort, helping identify where additional standardized surveillance would yield the greatest improvement in spatial information.

Several limitations should be considered. The underlying studies varied in sampling year, host composition, duration, protocols, and intensity; standardizing abundance by sampling effort improved comparability but could not remove all heterogeneity. Unequal original sample sizes may also affect district-level precision, which was addressed by explicitly quantifying posterior uncertainty. Environmental and livestock covariates represented broad spatial conditions rather than conditions matched to each sampling period, limiting temporal interpretation despite the use of long-term environmental summaries and harmonized livestock data. Finally, the analysis focused on tick abundance rather than pathogen transmission, so predictions were interpreted as indicators of tick abundance and surveillance need rather than direct measures of disease risk.

## 5. Conclusion

This study generated district-level Bayesian spatial predictions of the abundance of *R. microplus* and *H. anatolicum* across Punjab and KP using heterogeneous published surveillance data. The two species showed contrasting spatial and environmental patterns, with *R. microplus* exhibiting greater geographic heterogeneity and *H. anatolicum* showing relatively higher predicted abundance in southern Punjab. By combining exposure-adjusted observations with spatial prediction and posterior uncertainty, the analysis distinguishes areas supported by relatively informative evidence from districts where additional standardized surveillance would be most valuable. The framework provides a practical approach for extracting spatial information from fragmented veterinary surveillance data and could be extended to other vectors and data-sparse animal-health systems.

## Credit authorship contribution statement

**Abrar Hussain**: Conceptualization, Methodology, Data Curation, Data visualization, Formal analysis, Software, Writing an original draft, Review & editing; **Sabir Hussain**: Methodology, Review & editing; **Lelys Bravo de Guennic**: Methodology, Formal analysis, Review & editing; **Rebecca L. Smith**: Conceptualization, Review & editing, Supervision.

## Data availability

The relevant codes and data can be accessed here: https://github.com/abrarhussain95/PAK_tick

## Ethics statement

This study used aggregated data extracted from previously published studies and did not involve direct animal handling, experimentation, or collection of new biological samples. Therefore, institutional animal ethics approval was not required for the present analysis.

## Acknowledgments

We sincerely appreciate the efforts of all researchers who have documented tick occurrence records.

