## Supplementary material for "Bayesian spatial prediction of livestock tick abundance to support surveillance in data-sparse regions": Suplimentary Material

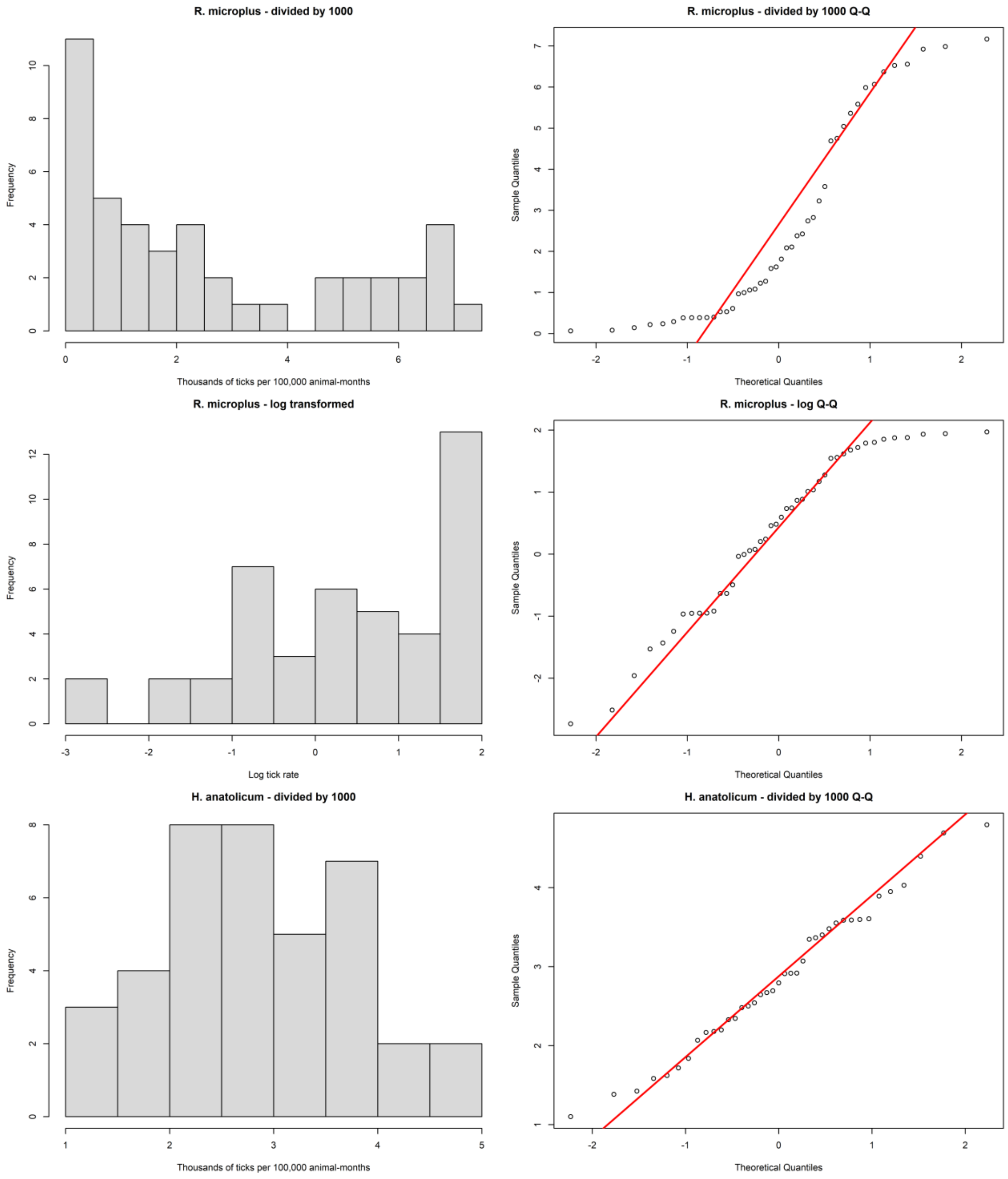


**Figure S1** Distributional assessment of species-specific exposure-adjusted tick abundance rates. Histograms and normal Q-Q plots are shown for the original *Rhipicephalus microplus* abundance rate, the natural-log-transformed *R. microplus* rate, and the original *Hyalomma anatolicum* abundance rate. The log transformation reduced the strong right-skewness of the *R. microplus* outcome, whereas the *H. anatolicum* rate was approximately Gaussian on its original scale.

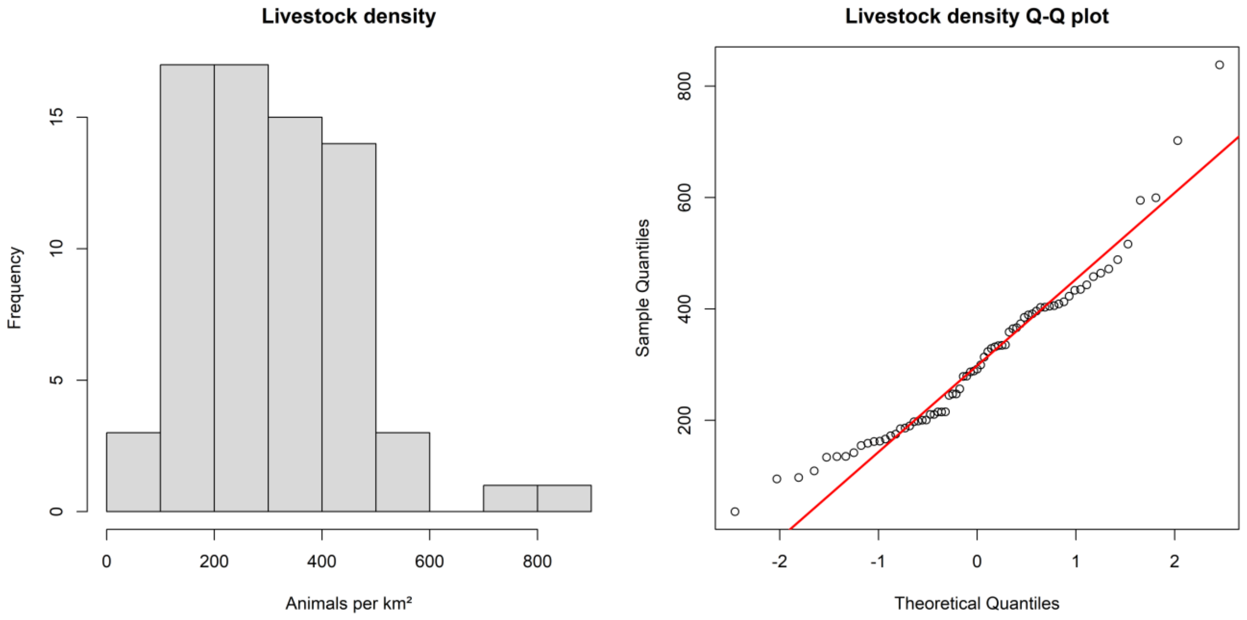


**Figure S2** Distribution of district-level livestock density. Histogram and normal Q-Q plot of livestock density (animals per km²) across study districts. Livestock density showed mild right-skewness, with most observations concentrated at moderate densities and a small number of high-density districts. The variable was standardized prior to inclusion in the regression models.

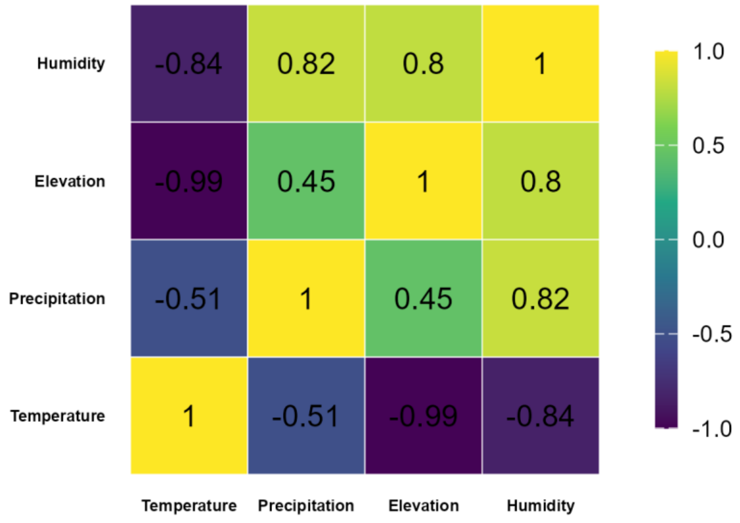


**Figure S3** Heatmap of pairwise Pearson correlation coefficients among district-level environmental covariates used in the analysis. Positive and negative values indicate the direction and strength of associations among mean temperature, precipitation, relative humidity, and elevation.

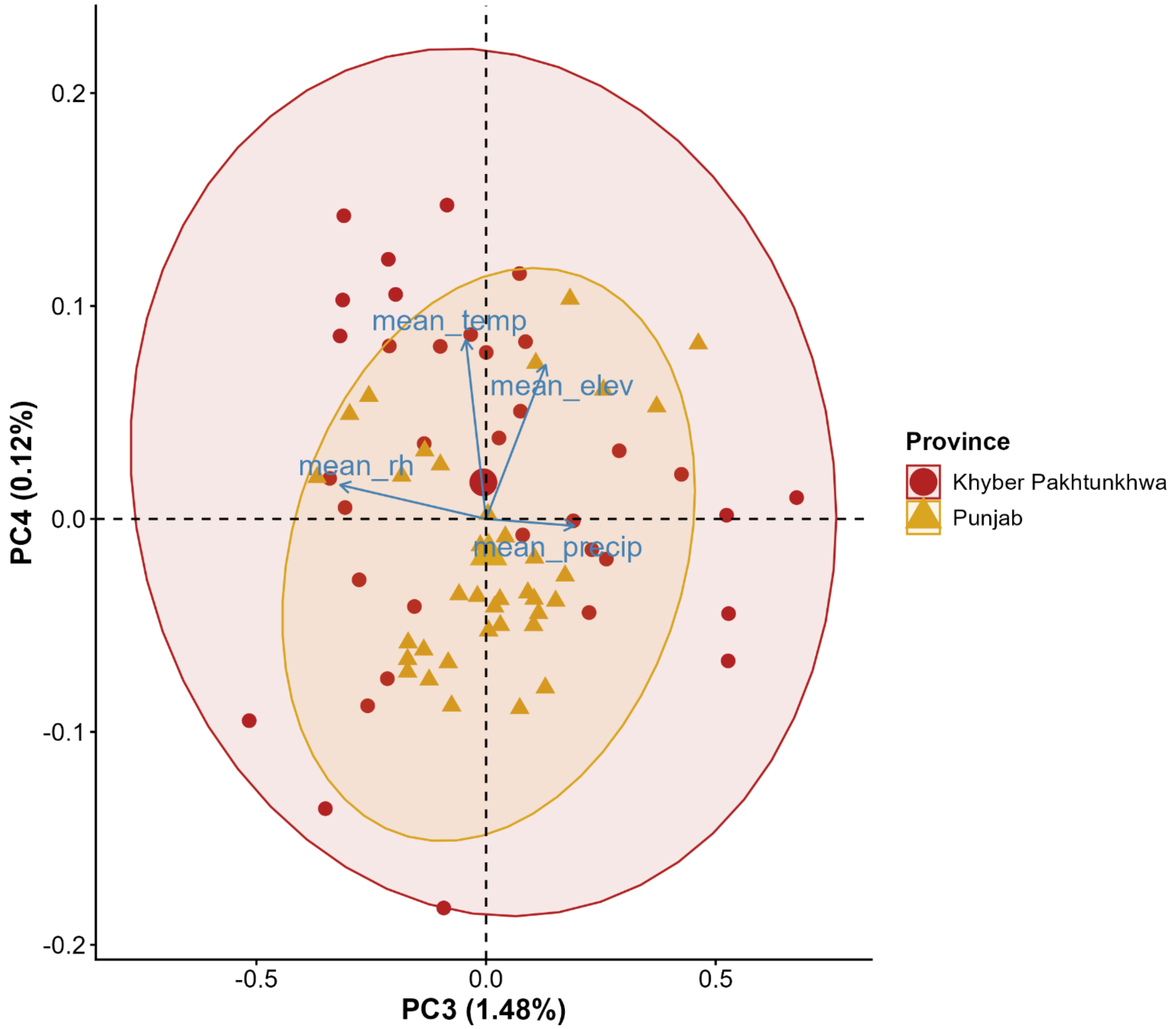


**Figure S4.** Principal component analysis biplot of PC3 and PC4 for district-level environmental covariates across Punjab and Khyber Pakhtunkhwa. The biplot shows district scores on PC3 and PC4, with vectors indicating the direction and relative contribution of mean temperature, precipitation, relative humidity, and elevation. Districts are distinguished by province, with ellipses indicating their distribution within each province.

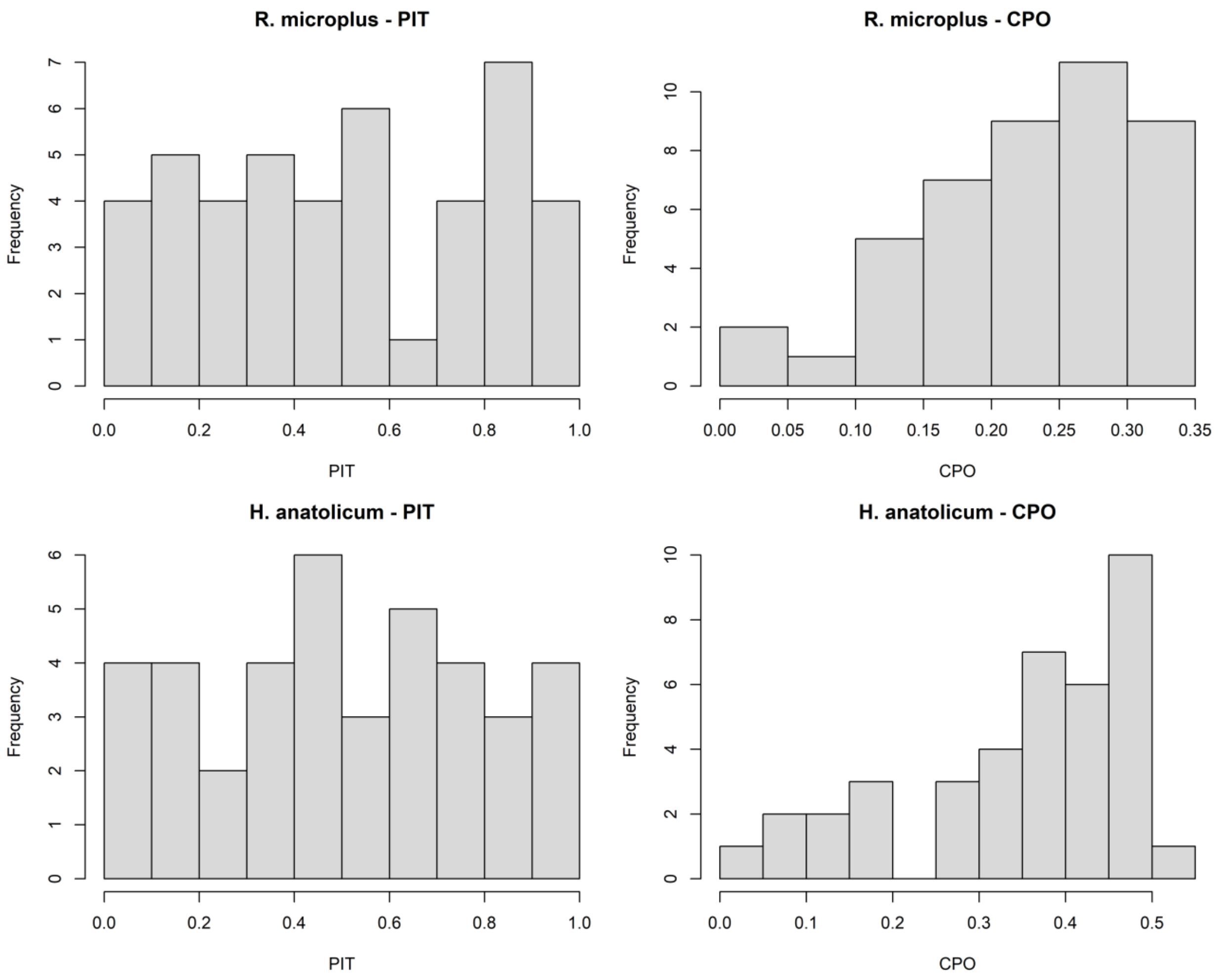


**Figure S5**. Conditional predictive ordinate (CPO) and probability integral transform (PIT) diagnostics for the final spatial BYM2 models of *Rhipicephalus microplus* and *Hyalomma anatolicum*. CPO values summarize leave-one-out predictive performance, while PIT distributions assess calibration of the posterior predictive distributions.

**Table S1.** Comparison of nonspatial Gaussian and spatial BYM2 models for *Rhipicephalus microplus* and *Hyalomma anatolicum* based on the Watanabe-Akaike Information Criterion (WAIC) and Deviance Information Criterion (DIC).

| Species | Model | DIC | WAIC |
|---|---|---|---|
| *R. microplus* | Nonspatial PC1-PC4 | 145.31 | 144.23 |
| *R. microplus* | Spatial BYM2 PC1-PC4 | 143.59 | 143.21 |
| *H. anatolicum* | Nonspatial PC1-PC4 | 96.28 | 96.07 |
| *H. anatolicum* | Spatial BYM2 PC1-PC4 | 90.97 | 92.88 |

**Table S2.** District-level leave-one-district-out validation results for *Rhipicephalus microplus* and *Hyalomma anatolicum*. Observed and posterior median predicted abundance are shown with corresponding 95% credible intervals (CrIs) and 95% predictive intervals (PIs). Coverage indicators denote whether the observed abundance fell within the respective interval.

| Species | District | Observed | Predicted | CrI_Lower95 | CrI_Upper95 | CrI_Covered95 |
|---|---|---|---|---|---|---|
| *R. microplus* | abbottabad | 2423 | 2213.465 | 255.8337 | 19457.34 | TRUE |
| *R. microplus* | mardan | 2087 | 5033.681 | 1530.103 | 17169.33 | TRUE |
| *R. microplus* | nowshera | 5045 | 1628.196 | 693.3243 | 3909.158 | FALSE |
| *R. microplus* | peshawar | 964 | 1334.888 | 457.0435 | 3993.642 | TRUE |
| *R. microplus* | shangla | 1813 | 1688.124 | 423.8945 | 6735.215 | TRUE |
| *R. microplus* | swabi | 5986 | 2610.548 | 940.0552 | 7466.367 | TRUE |
| *R. microplus* | swat | 6526 | 7611.393 | 1586.336 | 36362.12 | TRUE |
| *R. microplus* | tank | 6988 | 432.2517 | 43.45 | 4250.613 | FALSE |
| *R. microplus* | upper dir | 6922 | 5236.603 | 654.5166 | 41523.52 | TRUE |
| *R. microplus* | attock | 2107 | 989.0801 | 350.539 | 2795.474 | TRUE |
| *R. microplus* | bahawalnagar | 141 | 518.4595 | 185.0533 | 1463.758 | FALSE |
| *R. microplus* | bannu | 399 | 470.243 | 107.0573 | 2075.061 | TRUE |
| *R. microplus* | bahawalpur | 217 | 330.0639 | 89.54493 | 1195.718 | TRUE |
| *R. microplus* | bhakkar | 531 | 724.017 | 290.0054 | 1823.15 | TRUE |
| *R. microplus* | chakwal | 1081 | 884.0044 | 325.7573 | 2371.973 | TRUE |
| *R. microplus* | dera ghazi khan | 387 | 681.9741 | 196.2794 | 2398.921 | TRUE |
| *R. microplus* | faisalabad | 6559 | 1722.648 | 650.4604 | 4610.562 | FALSE |
| *R. microplus* | gujranwala | 3227 | 1735.512 | 633.8261 | 4291.09 | TRUE |
| *R. microplus* | gujrat | 996 | 1368.275 | 362.3318 | 5103.298 | TRUE |
| *R. microplus* | hafizabad | 1058 | 1850.086 | 760.1419 | 4298.883 | TRUE |
| *R. microplus* | jhang | 609 | 1312.403 | 554.5214 | 3132.695 | TRUE |
| *R. microplus* | jhelum | 2822 | 793.987 | 277.279 | 2202.72 | FALSE |
| *R. microplus* | batagram | 4751 | 4866.235 | 457.4796 | 51436.69 | TRUE |
| *R. microplus* | kasur | 4688 | 1353.128 | 461.832 | 3752.643 | FALSE |
| *R. microplus* | khanewal | 5584 | 1048.057 | 421.7391 | 2602.788 | FALSE |
| *R. microplus* | khushab | 2381 | 760.8084 | 309.0451 | 1853.217 | FALSE |
| *R. microplus* | lahore | 381 | 1414.28 | 542.8368 | 3510.881 | FALSE |

| | | | | | | |
|---|---|---|---|---|---|---|
| *R. microplus* | mandi bahauddin | 2740 | 1723.986 | 681.0457 | 4166.251 | TRUE |
| *R. microplus* | mianwali | 386 | 723.0279 | 286.022 | 1840.756 | TRUE |
| *R. microplus* | multan | 1226 | 1210.691 | 410.2805 | 3567.625 | TRUE |
| *R. microplus* | muzaffargarh | 288 | 1182.271 | 429.7662 | 3289.351 | FALSE |
| *R. microplus* | okara | 6066 | 1193.147 | 502.545 | 2825.22 | FALSE |
| *R. microplus* | rawalpindi | 385 | 1814.779 | 659.1413 | 5102.299 | FALSE |
| *R. microplus* | buner | 5363 | 1850.728 | 566.4807 | 6195.122 | TRUE |
| *R. microplus* | sahiwal | 6371 | 1059.537 | 406.182 | 2826.455 | FALSE |
| *R. microplus* | sargodha | 531 | 1694.694 | 722.1225 | 3970.577 | FALSE |
| *R. microplus* | sheikhupura | 65 | 1938.352 | 817.1137 | 4482.409 | FALSE |
| *R. microplus* | toba tek singh | 1620 | 1366.369 | 552.7826 | 3510.351 | TRUE |
| *R. microplus* | vehari | 239 | 1143.222 | 432.1632 | 3066.667 | FALSE |
| *H. anatolicum* | abbottabad | 2500 | 2293.429 | 790.9997 | 3711.129 | TRUE |
| *H. anatolicum* | bannu | 2671 | 1806.79 | 688.9079 | 2931.75 | TRUE |
| *H. anatolicum* | batagram | 1102 | 1194.818 | 536.326 | 2954.24 | TRUE |
| *H. anatolicum* | charsadda | 1584 | 2325.023 | 1228.02 | 3388.213 | TRUE |
| *H. anatolicum* | haripur | 1384 | 2567.452 | 1561.097 | 3510.584 | FALSE |
| *R. microplus* | charsadda | 1274 | 4256.972 | 1074.735 | 17247.17 | TRUE |
| *H. anatolicum* | karak | 2794 | 2545.973 | 1608.874 | 3485.851 | TRUE |
| *H. anatolicum* | kohat | 1838 | 2520.715 | 1662.892 | 3359.912 | TRUE |
| *H. anatolicum* | mansehra | 2345 | 2748.916 | 1443.905 | 4063.199 | TRUE |
| *H. anatolicum* | mardan | 2917 | 2588.472 | 1452.373 | 3640.874 | TRUE |
| *H. anatolicum* | peshawar | 2179 | 2043.612 | 1086.528 | 2930.598 | TRUE |
| *H. anatolicum* | shangla | 1717 | 1489.781 | 434.9257 | 2543.609 | TRUE |
| *H. anatolicum* | swat | 3347 | 2429.872 | 905.9225 | 3973.449 | TRUE |
| *H. anatolicum* | tank | 2911 | 3192.17 | 2275.984 | 4144.39 | TRUE |
| *H. anatolicum* | attock | 2067 | 2636.128 | 1742.211 | 3476.526 | TRUE |
| *H. anatolicum* | bahawalnagar | 2694 | 3406.468 | 2529.912 | 4306.024 | TRUE |
| *R. microplus* | karak | 81 | 599.4235 | 187.3835 | 1937.046 | FALSE |
| *H. anatolicum* | bahawalpur | 3596 | 3726.962 | 2754.905 | 4691.166 | TRUE |
| *H. anatolicum* | bhakkar | 4693 | 3169.931 | 2292.622 | 4028.156 | FALSE |
| *H. anatolicum* | chakwal | 2542 | 2618.007 | 1766.322 | 3455.806 | TRUE |
| *H. anatolicum* | dera ghazi khan | 3606 | 3905.965 | 2913.777 | 4900.94 | TRUE |
| *H. anatolicum* | gujranwala | 2199 | 3075.65 | 2094.134 | 4137.094 | TRUE |
| *H. anatolicum* | gujrat | 2919 | 2501.436 | 1552.695 | 3517.628 | TRUE |
| *H. anatolicum* | jhelum | 2481 | 2531.401 | 1680.792 | 3411.658 | TRUE |
| *H. anatolicum* | kasur | 3587 | 2894.365 | 1957.595 | 3797.164 | TRUE |
| *H. anatolicum* | khanewal | 4030 | 3448.298 | 2582.339 | 4281.381 | TRUE |
| *H. anatolicum* | khushab | 3071 | 3232.641 | 2458.235 | 4034.199 | TRUE |
| *R. microplus* | kohat | 1582 | 515.7025 | 182.1587 | 1460.686 | FALSE |
| *H. anatolicum* | lahore | 1622 | 2968.167 | 2201.222 | 3788.638 | FALSE |
| *H. anatolicum* | mandi bahauddin | 3365 | 2733.826 | 1976.346 | 3542.928 | TRUE |
| *H. anatolicum* | mianwali | 2167 | 2808.39 | 2015.595 | 3600.237 | TRUE |
| *H. anatolicum* | multan | 3479 | 3709.145 | 2787.867 | 4606.74 | TRUE |
| *H. anatolicum* | muzaffargarh | 2644 | 3827.843 | 2968.275 | 4705.614 | FALSE |
| *H. anatolicum* | okara | 4398 | 3216.003 | 2320.672 | 4067.254 | FALSE |
| *H. anatolicum* | rahim yar khan | 4795 | 3477.037 | 2471.68 | 4435.8 | FALSE |

| | | | | | | |
|---|---|---|---|---|---|---|
| *H. anatolicum* | rawalpindi | 2328 | 2492.069 | 1610.332 | 3308.379 | TRUE |
| *H. anatolicum* | sahiwal | 1425 | 3577.772 | 2835.11 | 4324.272 | FALSE |
| *H. anatolicum* | sargodha | 3402 | 3096.41 | 2329.678 | 3898.444 | TRUE |
| *R. microplus* | malakand | 7168 | 2732.726 | 891.2719 | 8454.729 | TRUE |
| *H. anatolicum* | sheikhupura | 3589 | 3080.466 | 2309.471 | 3911.074 | TRUE |
| *H. anatolicum* | sialkot | 3894 | 2200.096 | 1284.86 | 3155.606 | FALSE |
| *H. anatolicum* | toba tek singh | 3553 | 3411.98 | 2551.656 | 4231.057 | TRUE |
| *H. anatolicum* | vehari | 3950 | 3524.054 | 2641.067 | 4352.99 | TRUE |
| *R. microplus* | mansehra | 3580 | 8811.54 | 2109.961 | 36549.73 | TRUE |
